# Asymmetric Cross-Reactivity of Nuclear Receptors Reveals an Evolutionary Buffer Between Estrogen and Androgen Signaling

**DOI:** 10.64898/2025.12.18.695200

**Authors:** Shintaro Yamazaki, Akhilesh B. Reddy

## Abstract

A comprehensive all-by-all receptor ligand affinity screen using Boltz-2, a deep learning framework for protein-ligand interaction prediction, reveals a previously unrecognized asymmetry in steroid hormone receptor binding. Using systematic *in silico* affinity prediction, we show that estradiol binds the androgen receptor with higher predicted affinity than testosterone and also displays strong affinity for multiple related steroid hormone receptors, whereas the reciprocal interaction – binding of non-aromatized steroids to estrogen receptors – is not observed. Structural modeling demonstrates that estradiol and testosterone occupy the same canonical ligand-binding pocket within the androgen receptor, indicating a conserved steroid-recognition architecture rather than a specialized binding mode. Analysis of a clinically relevant androgen receptor mutation shows modest and broadly distributed stabilization of ligand engagement, consistent with tuning of a pre-existing estradiol-compatible interaction rather than generation of a novel binding mechanism. Reconstruction of ancestral steroid receptors indicates that estradiol maintains high predicted affinity across both ancestral and modern receptors, while other steroids progressively diversify in their receptor preferences following lineage expansion. Together, these results support an evolutionary model in which estradiol represents an early steroid ligand, with younger receptors retaining ancestral estrogen compatibility while evolving specificity for upstream steroid hormones. Functionally, this asymmetric architecture provides a mechanism by which estradiol may modulate androgen receptor signaling under physiological conditions and may contribute to altered receptor activation in pathological contexts such as advanced prostate cancer. These findings define a coherent biochemical and evolutionary framework for estradiol cross-reactivity and highlight the estradiol-androgen receptor interface as a potential therapeutic target.

## Introduction

Nuclear receptors (NRs) constitute a large family of ligand-regulated transcription factors that control diverse endocrine, metabolic and developmental processes. Despite extensive study, NR interaction networks remain incompletely characterized, in part because many receptors are classified as orphan receptors lacking identified endogenous ligands (Delfosse *et al*. 2015). Of the 48 human nuclear receptors, approximately half have no firmly established physiological ligand (Delfosse *et al*. 2015; Frigo *et al*. 2021). Even among receptors with known cognate hormones, potential secondary ligands and cross-reactive interactions are not comprehensively mapped, suggesting that functionally relevant crosstalk within the NR superfamily may remain underappreciated (De Bosscher *et al*. 2020).

Among nuclear receptors, the NR3 subfamily (comprising steroid hormone receptors) provide a biologically and clinically-relevant system in which to explore ligand cross-interactions. These receptors bind hormones derived from a common cholesterol-based biosynthetic cascade and share a conserved structural framework (Eick and Thornton 2011; Thornton 2001). The prevailing paradigm assumes one-to-one ligand-receptor specificity, with ERα/β responding to estrogens and AR responding to androgens. However, multiple empirical observations challenge this simple model. In prostate tissue, estradiol has been reported to suppress AR signaling and promote AR destabilization under physiological conditions (Woodham *et al*. 2003), whereas in advanced prostate cancer estradiol can activate AR signaling in the presence of permissive coactivator environments or ligand-binding domain mutations (Elo *et al*. 1995; Yeh *et al*. 1998). These context-dependent effects remain mechanistically unresolved.

Evolutionary analyses have provided important insights into the origins of steroid hormone receptors. The common ancestor of modern steroid receptors is thought to have been estrogen-responsive, exhibiting high affinity for aromatized ligands such as estradiol. Subsequent gene duplication and diversification events gave rise to the NR3 subfamily, including estrogen receptors (ERα and ERβ), androgen receptor (AR), progesterone receptor (PR), glucocorticoid receptor (GR), and mineralocorticoid receptor (MR) (Baker 2019; Eick and Thornton 2011). While these receptors evolved specificity for distinct pregnenolone-derived steroids, such as testosterone, progesterone, cortisol, and aldosterone, they retained a conserved ligand-binding scaffold inherited from an estrogen-sensitive ancestor (Thornton 2001). This shared structural ancestry raises the possibility that modern NR3 receptors may preserve latent compatibility with estrogens, despite their apparent ligand specificity.

Systematic investigation of NR-ligand interactions has traditionally relied on targeted biochemical assays, which are low throughput and typically restricted to expected receptor-ligand pairs. Recent advances in computational modeling now offer an alternative strategy. In particular, Boltz-2, a deep learning based protein-ligand co-folding framework, enables high-throughput prediction of receptor-ligand structures and binding affinities with accuracy approaching that of more computationally intensive free-energy methods (Passaro *et al*. 2025). This approach makes it feasible to perform unbiased, all-by-all surveys of NR-ligand interactions at a scale, not achievable experimentally with wet-lab assays alone.

In this study, we used an unbiased computational framework to systematically map ligand affinity landscapes across the human nuclear receptor superfamily. By integrating all-by-all affinity prediction, structural modeling, receptor mutation analysis, and ancestral receptor reconstruction, we aimed to determine whether conserved evolutionary features of NR3 receptors contribute to asymmetric ligand cross-reactivity. Our results identify estradiol as a broadly compatible ligand across the NR3 lineage and support an evolutionarily conserved mechanism by which estrogen signaling could modulate other receptor activity under both physiological and pathological conditions.

## Materials and methods

### Nuclear receptor sequence selection

Human nuclear receptors (48 receptors in total; Table 1) were included in an unbiased computational screen. Full-length canonical protein sequences were obtained from UniProt and used without truncation. To examine the impact of receptor modification on ligand affinity landscapes, the androgen receptor T877A mutation was modeled *in silico*. This mutation was selected based on its established role in altering steroid responsiveness in prostate cancer (Sun *et al*. 2006). The mutant AR sequence was generated by single-residue substitution within the full-length receptor and subjected to the same all-by-all ligand screening pipeline as wild-type AR. Reconstructed ancestral nuclear receptor sequences (AncSR1 and AncSR2) were obtained from previously published phylogenetic analyses of the NR3 lineage (Eick *et al*. 2012). These sequences were derived using maximum-likelihood methods and experimentally validated in prior work (Eick *et al*. 2012). Ancestral receptors were treated equivalently to extant receptors in all computational analyses.

**Table 1.** List of the 48 canonical human nuclear hormone receptors included in the all-by-all affinity screen, with NR classifications, gene symbols and known functions.

### Ligand library construction

This comprehensive collection includes all major endogenous nuclear hormone receptor ligands, including steroid hormones, thyroid hormones, vitamin D metabolites, retinoids, fatty acids, and bile acids (Table 2). All ligand identities were verified through systematic PubChem searches, and all PubChem compound identifiers (CIDs) were cross-checked. Ligands were not pre-assigned to cognate receptors.

**Table 2.** List of endogenous ligands used in the study.

### Ligand-receptor affinity prediction using Boltz-2

Ligand-receptor interactions were predicted using Boltz-2, a deep learning modeling framework trained on experimentally determined receptor-ligand structural data (Passaro *et al*. 2025). Data were generated on a compute node with 8x Tesla P40 Nvidia GPUs (24GB each), 512GB RAM and 4TB SSD. For each receptor-ligand pair, Boltz-2 predicts an IC₅₀ value (nM) reflecting binding strength and an associated confidence score reflecting model certainty. All receptors were screened against the complete ligand library, generating an unbiased all-by-all interaction matrix. Confidence scores were used only for post hoc filtering and visualization, with specific thresholds indicated in the corresponding figure legends.

### Molecular docking and structural comparison

To independently evaluate ligand binding modes, estradiol and testosterone were docked into the androgen receptor using AutoDock Vina (Trott and Olson 2010). Docking was performed using a cubic search grid (40 × 40 × 40 Å) centered on the receptor center of mass, encompassing the canonical AR ligand-binding cavity defined in prior structural studies (Sack *et al*. 2001). For each ligand, the highest-scoring pose located within the canonical binding pocket was selected. Structural visualization and superposition were performed using ChimeraX (Pettersen *et al*. 2021).

### Evolutionary integration and comparative analysis

Predicted ligand affinities for extant and ancestral receptors were interpreted in the context of established NR3 phylogeny and steroid biosynthetic pathways. Divergence timing and ancestral functional states were based on prior evolutionary analyses (Eick *et al*. 2012). Steroid biosynthetic relationships were overlaid conceptually to relate predicted receptor affinities to the emergence of enzymatic steps such as aromatization by CYP19A1 (Santen *et al*. 2009).

### Data visualization and analysis

All figures were generated in R (version 4.4.0) using ggplot2 (version 4.0.0). Bar plots display predicted log₁₀(IC₅₀) values ordered by affinity, with color intensity representing prediction confidence. Scatter plots comparing wild-type and mutant AR affinities were plotted on matched log scales, with the diagonal indicating equal predicted affinity.

## Results

### Comprehensive all-by-all affinity screen reveals asymmetric cross-reactivity in NR3 subfamily

To systematically map potential cross-reactivity across nuclear receptors, we performed an exhaustive all-by-all *in silico* affinity screen using full-length sequences of all 48 human nuclear receptors spanning NR0-NR6 and a curated library of 54 endogenous nuclear receptor ligands, including steroid hormones and intermediates, thyroid hormones, vitamin D metabolites, retinoids, fatty acids, and bile acids. Predicted ligand-receptor affinities and confidence scores were generated using Boltz-2 (Passaro *et al*. 2025), yielding a complete interaction matrix across the human nuclear receptor repertoire (Supplementary File 1).

As an internal benchmark, the screen recovered established cognate ligand preferences with appropriate rank ordering. For example, ERα/ERβ showed strongest predicted binding to 17β-estradiol (E2), followed by estrone (E1) and estriol (E3), consistent with known relative estrogenic potency (Kuiper *et al*. 1997) (Figure 1A,B). Vitamin D receptor (VDR) preferentially bound calcitriol (1,25-dihydroxyvitamin D₃) relative to calcidiol (25-hydroxyvitamin D₃) and vitamin D₂, consistent with its established hormonal hierarchy (Reichel *et al*. 1989; Teske *et al*. 2016) (Figure 1C). Likewise, thyroid hormone receptors (TRα/TRβ) preferentially bound T3 over the prohormone T4, consistent with established TR selectivity (Oppenheimer and Schwartz 1997; Wejaphikul *et al*. 2019) (Figure 1D,E). Notably, neither VDR nor TR displayed high-confidence binding to steroid hormones, including estradiol or androgens.

**Figure 1.**
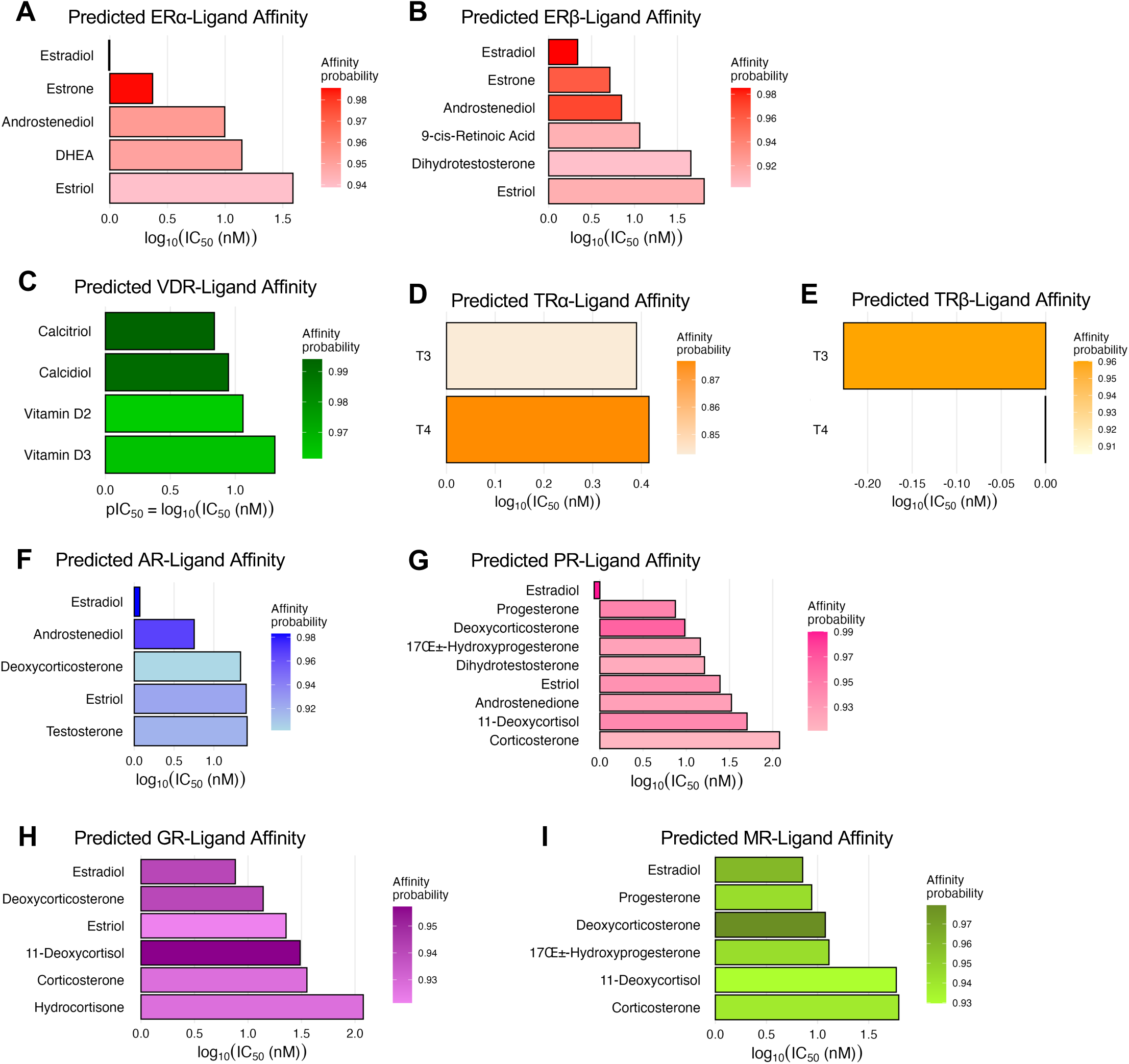
Boltz-2 predicts asymmetric estradiol affinities across the NR3 subfamily. Bar plots show predicted ligand affinities (x-axis: log₁₀(IC₅₀ [nM])) for each receptor. Lower values indicate stronger prediction binding. Color intensity represents Boltz-2 prediction confidence (scale bar at right), with darker bars indicating higher model certainty (maximum = 1.00). Only ligands with prediction confidence >0.90 are displayed (except for TRα, which had no ligands exceeding this threshold). IC₅₀ and confidence scores are derived from the Boltz-2 model, a deep learning framework trained on receptor-ligand structural data to jointly predict binding strength and interaction likelihood. (A) ERα binds estradiol with highest predicted affinity, followed by estrone and androstene-derived metabolites. (B) ERβ shows a similar rank order, with estradiol top-ranked, and modest binding predicted for 9-cis-retinoic acid and DHT. (C) VDR shows high affinity and confidence for calcitriol, followed by calcidiol and vitamin D2/D3, matching known biology. (D) TRα ranks T3 over T4, but neither ligand exceeds the confidence threshold (>0.90). (E) TRβ shows higher-confidence binding for T3 over T4, consistent with TRβ’s known ligand preference. (F) AR is predicted to bind estradiol with high affinity and high model confidence, exceeding even androstene-based ligands. (G) PR shows top binding to estradiol and progesterone, with notable affinity for multiple steroid intermediates including DOC and 17α-hydroxyprogesterone. (H) GR binds estradiol comparably to glucocorticoids such as cortisol, corticosterone, and their precursors. (I) MR shows high-confidence affinity for estradiol, alongside known mineralocorticoids like aldosterone precursors.

Beyond these expected interactions, the global screen revealed an unexpected and pronounced asymmetry in cross-reactivity within NR3. E2 was predicted to bind with high affinity not only to ERα/ERβ, but also to multiple 3-ketosteroid receptors, AR, PR, GR, and MR, ranking among the strongest predicted ligands for these receptors beyond their cognate ligands (Figure 1F–I). In contrast, the reciprocal was not observed: canonical ligands for AR/PR/GR/MR (e.g. testosterone/DHT, progesterone, cortisol, aldosterone) did not show meaningful predicted binding to ERα or ERβ at comparable confidence thresholds (Figure 1A,B). Thus, the affinity landscape appears to be directional – estradiol engages multiple steroid receptors, whereas non-aromatized steroids remain excluded from estrogen receptors. Given our results from cognate binding preferences above, the observed estradiol cross-reactivity is not a general artifact of ligand promiscuity in the model, but a feature specific to the NR3 steroid receptor lineage.

### Estradiol binds AR with high predicted affinity, consistent with prior biochemical evidence

Among the cross-reactive interactions, the E2-AR pair was particularly notable. The screen predicted sub-nanomolar to low-nanomolar binding of E2 to AR, exceeding the predicted affinity of testosterone under the same analysis conditions (Figure 1F). In the AR-focused comparison, E2 was predicted at approximately ∼1.2 nM versus ∼26.2 nM for testosterone (∼20-fold difference) (Figure 1F), while testosterone showed negligible predicted affinity for ERα/β (Figure 1A,B).

This prediction is consistent with prior experimental studies showing that AR can directly bind estradiol (Veldscholte *et al*. 1992). In prostate cell contexts, Yeh et al. demonstrated that estradiol can drive AR transcriptional activity in the presence of the AR coactivator ARA70, implying productive occupancy of the AR ligand-binding pocket under permissive cofactor conditions (Yeh *et al*. 1998). Competitive binding experiments have reported E2-AR binding IC₅₀ values in the low-nanomolar range in relevant assay settings (Yeh *et al*. 1998). Conversely, androgen binding to ERα is extremely weak relative to estradiol (Kuiper *et al*. 1997; Rider *et al*. 2009), consistent with the asymmetry observed in our screen.

Extending beyond AR, E2 also showed high predicted affinity for PR, GR, and MR (Figure 1G-I), supporting the interpretation that estradiol displays an unusual degree of compatibility across 3-ketosteroid receptors, whereas those receptors’ principal ligands do not reciprocally engage ERα/β (Figure 1A,B; Figure 1F-I).

### Estradiol engages the androgen receptor through a conserved steroid-recognition pocket

Importantly, this preference did not arise from an alternative or aberrant binding mode. Molecular docking demonstrated that estradiol and testosterone occupy the same canonical AR ligand-binding pocket (Nadal *et al*. 2017), with highly overlapping poses (Figure 2A). The shared occupancy of this pocket indicates a conserved steroid-recognition architecture in AR, capable of accommodating both aromatized and non-aromatized steroids. This architectural compatibility explains how estradiol can bind AR with high affinity despite lacking classical androgenic features. Critically, the docking results argue against a specialized or mutation-dependent binding mode for estradiol; instead, estradiol exploits an existing pocket whose geometry predates strict androgen specificity and was later refined for non-aromatized steroids. This interpretation is consistent with evolutionary reconstructions indicating that modern NR3 ligand-binding pockets derive from an estrogen-sensitive ancestral architecture that was subsequently tuned – but not replaced – during specialization for non-aromatized steroids (Thornton 2001).

**Figure 2.**
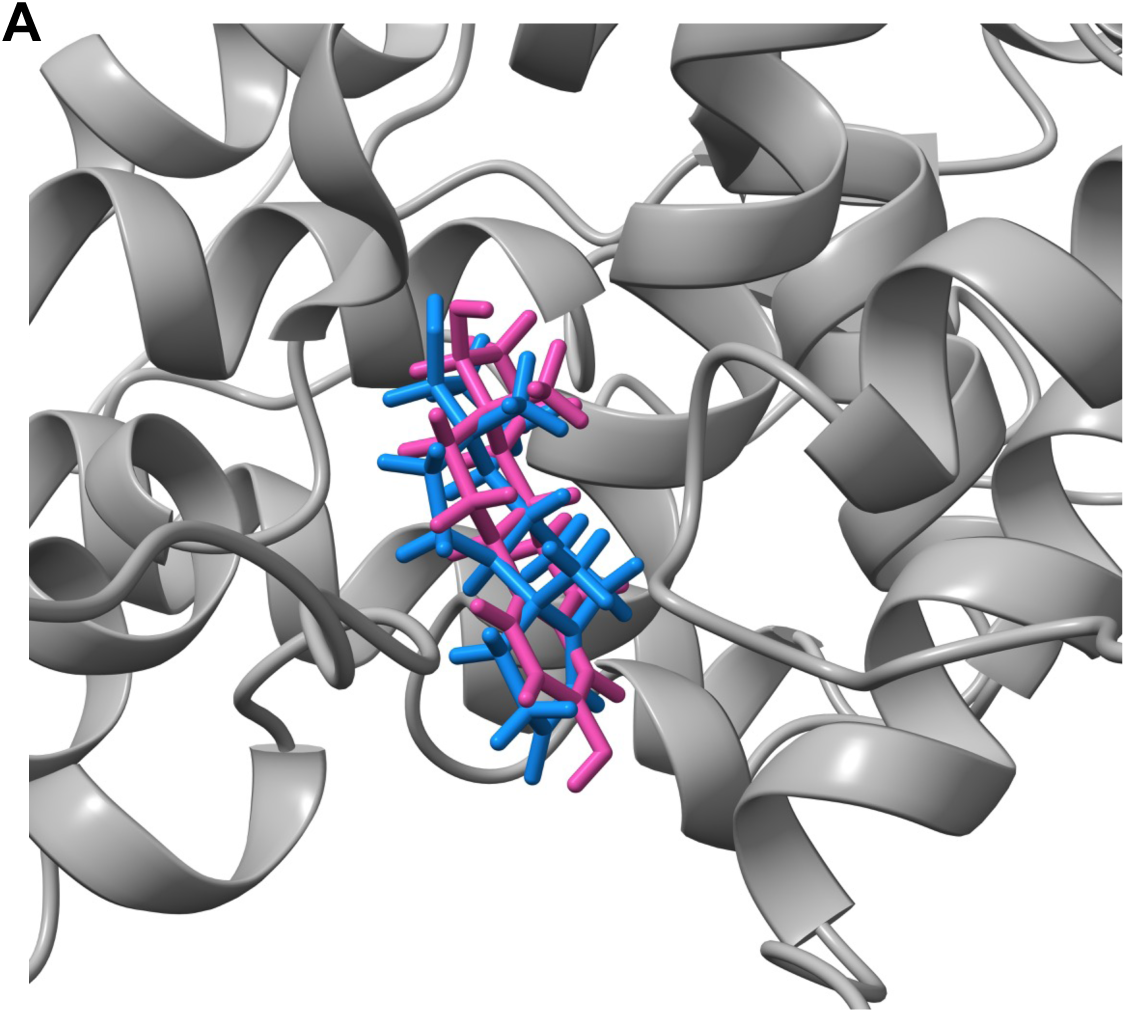
Estradiol and testosterone occupy a shared binding pocket in the androgen receptor. (A) Structural superposition of estradiol (magenta) and testosterone (blue) docked within the ligand-binding domain (LBD) of the androgen receptor (AR), shown as a gray ribbon representation. Both ligands adopt highly overlapping poses within the canonical AR binding cavity, indicating that estradiol is accommodated without a distinct binding mode. Docking was performed using AutoDock Vina with a blind search grid encompassing the AR LBD (40 × 40 × 40 Å) centered near the receptor center of mass, and the highest-scoring pose located within the canonical pocket was selected for analysis. Pose visualization was performed in ChimeraX. The shared occupancy of this pocket suggests a conserved steroid-recognition architecture in AR underlying asymmetric ligand affinity.

### AR T877A amplifies pre-existing estradiol compatibility rather than creating a novel interaction

To determine whether disease-associated AR mutations create new estradiol responsiveness or instead repurpose an existing interaction, we analyzed the T877A AR mutant, a clinically-relevant variant associated with ligand promiscuity in prostate cancer (Han *et al*. 2005; Veldscholte *et al*. 1992). Across the ligand panel, T877A produced modest, broadly distributed affinity gains, rather than selectively favoring a new ligand class (Figure 1F; Figure 3A).

**Figure 3.**
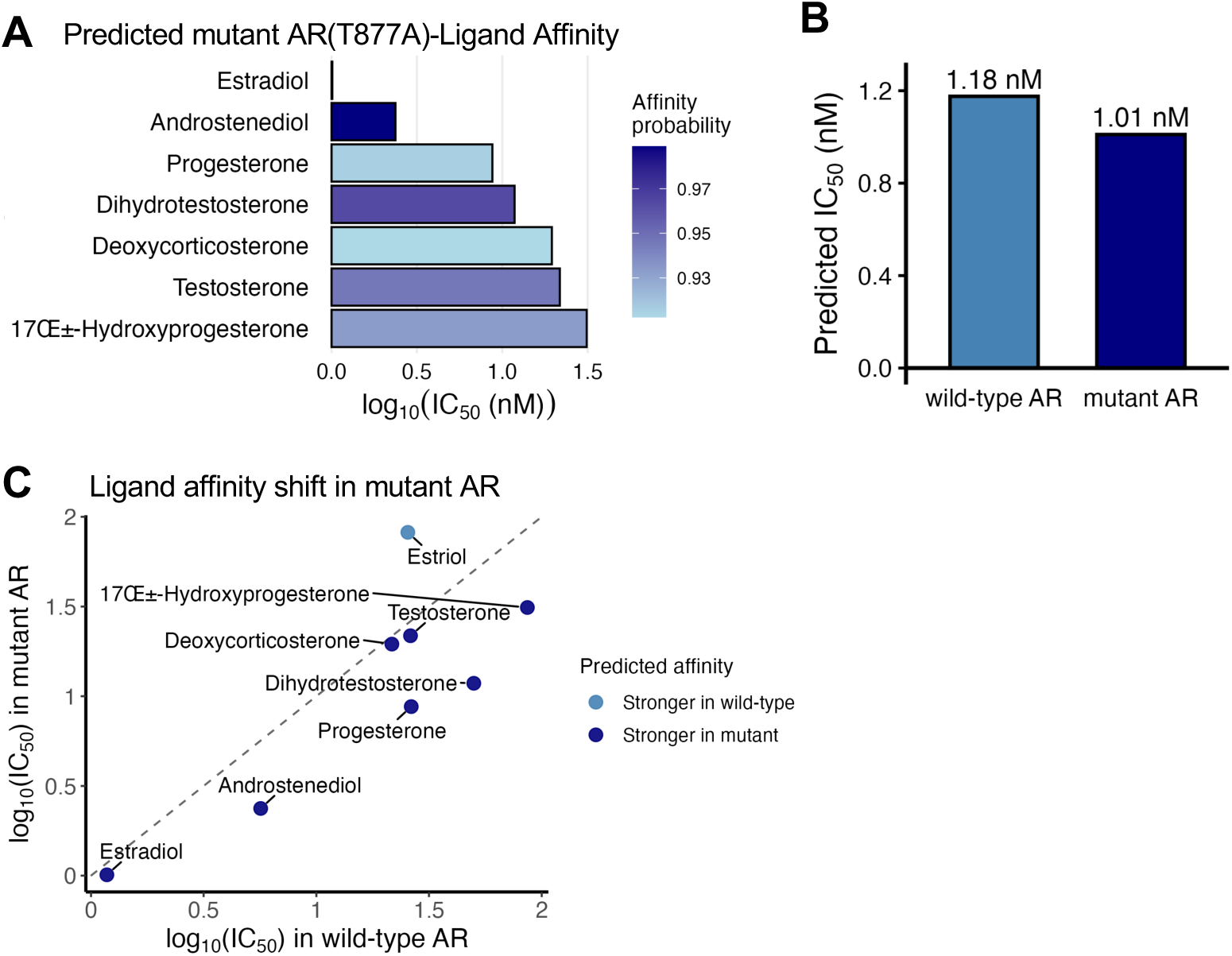
AR modification repurposes estradiol’s strong basal affinity for activation. (A) Predicted mutant AR (T877A)–ligand affinity: Boltz-2 modeling identifies multiple steroids with high predicted affinity and confidence (probability ≥0.93) toward the T877A mutant, including estradiol, androstenediol, progesterone, dihydrotestosterone, deoxycorticosterone, testosterone, and 17α-hydroxyprogesterone. Across the ligand panel, T877A produces modest and broadly distributed affinity gains, indicating that the mutation does not create a new high-affinity interaction but tunes the existing steroid-binding landscape. (B) Predicted estradiol affinity: Boltz-2 estimates similar IC₅₀ for estradiol binding to wild-type and T877A AR (predicted IC₅₀ ≈ 1.18 nM and 1.01 nM), consistent with estradiol retaining substantial AR compatibility regardless of mutation. Estradiol already shows strong basal compatibility with AR, and the mutation slightly enhances this pre-existing affinity, consistent with a model in which T877A repurposes ancestral estradiol binding for receptor stabilization rather than inventing a novel binding mode. (C) Affinity shift mutant AR: Scatterplot comparing wild-type and mutant AR predicted IC₅₀ (nM) values (log₁₀). Only ligands with high prediction confidence (probability ≥0.9) in either wild-type or mutant AR are shown. Points above the diagonal indicate ligands with weaker predicted affinity in the mutant; points below indicate stronger affinity in the mutant. Labels mark ligands with the largest predicted shifts.

Notably, estradiol remained the strongest-affinity ligand for AR even in the mutant context (Figure 3A), with predicted IC₅₀ values remaining similar between wild-type and mutant AR (Figure 3B). This indicates that estradiol already possesses strong basal compatibility with AR and that T877A slightly enhances a pre-existing interaction rather than inventing a new binding mode.

The pattern of affinity shifts across ligands supports this interpretation. T877A does not uniquely elevate affinity for a specific ligand class. Instead, it subtly stabilizes ligand engagement across the steroid spectrum (Figure 3C). Together with the shared-pocket docking results, this suggests that the mutation primarily tunes receptor conformational stability and coactivator permissiveness, rather than fundamentally altering ligand recognition or binding specificity.

### Ancestral receptor analysis indicates estradiol as the evolutionarily oldest NR3 ligand

To place the observed asymmetry in evolutionary context, we examined predicted affinities for reconstructed ancestral receptors AncSR1 and AncSR2 (Eick *et al*. 2012) (Figure 4A). The ancestral affinity profiles reveal a clear pattern: estradiol maintains consistently high predicted affinity across both ancestral (Figure 4B,C) and modern receptors (Figure 1A-B,F-I), whereas other steroids display progressively diversified affinity profiles following NR3 lineage expansion (Figure 4D).

**Figure 4.**
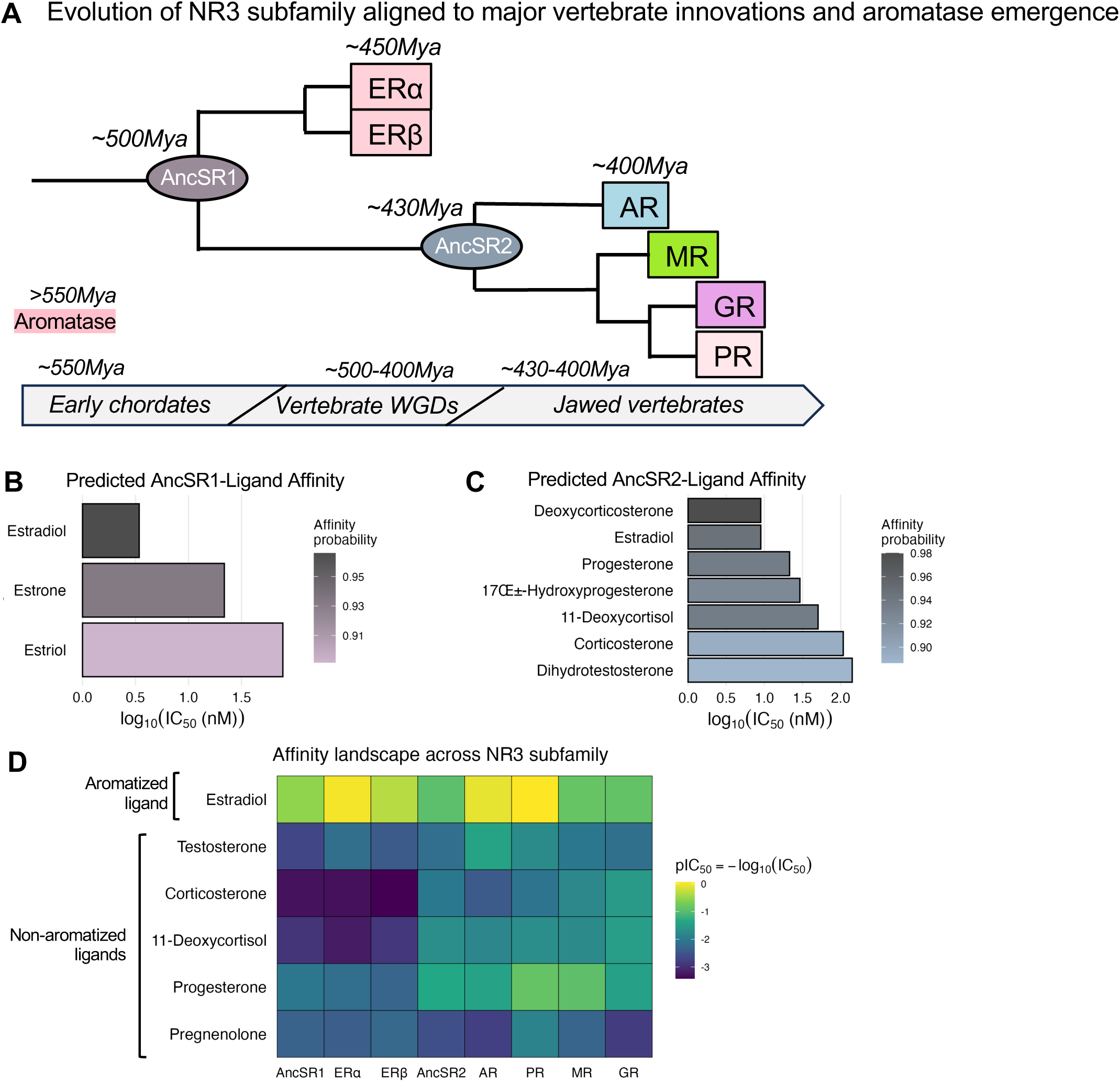
Ancestral receptor analysis reveals stepwise acquisition of non-aromatized steroid affinity across the NR3 lineage. (A) Phylogenetic relationships among NR3 receptors are shown with reconstructed ancestral nodes explicitly indicated. AncSR1 (∼500 Mya) represents the earliest estrogen-responsive ancestral steroid receptor, arising prior to vertebrate diversification. A subsequent duplication during early vertebrate whole-genome duplication events (WGDs) produced ERα and ERβ (∼450 Mya), both of which retained strict specificity for aromatized estrogens. A later divergence generated AncSR2 (∼430–400 Mya), the common ancestor of AR, PR, GR, and MR, coincident with the emergence of jawed vertebrates. This node marks the evolutionary transition at which responsiveness to non-aromatized, pregnenolone-derived steroids began to emerge, while ancestral estradiol compatibility was retained. (B, C) Predicted ligand affinities for AncSR1 and AncSR2 (probability≥0.874), respectively. AncSR1 shows strong and selective affinity for aromatized estrogens, consistent with an ancestral estrogen receptor. AncSR2 displays broader affinity across both aromatized and non-aromatized steroids, consistent with a transitional receptor that had begun to diversify ligand recognition while retaining ancestral compatibility with estradiol. (D) Affinity landscape across ancestral and modern NR3 receptors for key nodes of the steroidogenic cascade. Tile color represents predicted binding affinity expressed as *pIC₅₀ = −log₁₀(IC₅₀)*, with lighter (yellow) colors indicating stronger predicted affinity and darker colors indicating weaker affinity. Ligands are ordered according to their biosynthetic hierarchy (pregnenolone → progesterone → corticosteroids/androgens → estradiol), highlighting the alignment between receptor evolution and steroid metabolism. The heatmap reveals that estradiol maintains consistently high predicted affinity across ancestral and modern receptors, whereas other steroids show progressively diversified affinity profiles following NR3 lineage expansion. This pattern supports an evolutionary model in which modern receptors retain ancestral estradiol compatibility while diversifying specificity for downstream steroids.

AncSR1, representing the earliest reconstructed steroid receptor, shows strong preference for estradiol and limited responsiveness to non-aromatized steroids (Figure 4B). AncSR2 begins to acquire compatibility with non-aromatized ligands, but estradiol remains a high-affinity ligand (Figure 4C). In modern receptors, specificity for testosterone, progesterone, and corticosteroids increases, yet ancestral estradiol compatibility is retained (Figure 4D). Importantly, the emergence of aromatase (CYP19A1) predates the diversification of modern NR3 receptors (Santen *et al*. 2009) (Figure 4A), establishing estradiol as an ancient, pre-existing ligand in early vertebrates. Subsequent nuclear receptor evolution therefore occurred in the context of an already available estrogenic end-product, rather than requiring *de novo* ligand invention.

This evolutionary trajectory explains the asymmetric cross-reactivity observed in extant receptors. Although estradiol is the terminal product of steroid biosynthesis, it is the evolutionarily oldest ligand in the NR3 lineage. As the NR3 lineage expanded, younger receptors progressively diversified to recognize upstream non-aromatized steroids in the biosynthetic cascade, while retaining ancestral sensitivity to the downstream end-product estradiol (Figure 4A,D). In contrast, ERα/β, already optimized for estradiol recognition, never evolved responsiveness to non-aromatized steroids, reinforcing an asymmetric specialization.

Together, these data indicate that modern AR and related NR3 receptors do not appear to acquire estradiol binding adventitiously, but instead retain and repurpose an ancestral estrogen-compatible architecture while evolving specificity for newer steroid ligands.

## Discussion

### Asymmetric estradiol cross-reactivity as an organizing principle of NR3 signaling

A central finding of this study is that estradiol (E2) exhibits strong predicted affinity across multiple NR3 steroid receptors – including AR, PR, GR, and MR – whereas the converse is not observed: canonical non-aromatized NR3 ligands show minimal predicted affinity for ERα/β. This one-way cross-reactivity provides a coherent biochemical explanation for long-standing context-dependent observations in steroid endocrinology, particularly the paradox in which estradiol represses AR activity in healthy tissues yet can activate AR signaling in malignancy. Rather than representing non-specific promiscuity, the directionality suggests an inherited architectural constraint within the NR3 lineage – an “estrogen-first” compatibility retained in younger receptors that later specialized to bind upstream steroids.

### A two-state model for estradiol action on AR: buffering versus pathological repurposing

Our results support a two-state framework in which the functional outcome of E2 engagement of AR depends on receptor state and cellular context. In a physiological “buffering” state (Figure 5A), estradiol binds AR but typically fails to generate robust transcriptional output, consistent with prior evidence that E2-bound AR is inefficient at coactivator recruitment and downstream activation (Yeh *et al*. 1998). In this setting, estradiol can function as a brake on androgen signaling, supported by reports that estradiol promotes AR destabilization and proteasomal degradation (Woodham *et al*. 2003) and that developmental estradiol exposure can durably reduce AR protein in prostate tissue (Roberts *et al*. 2000; Woodham *et al*. 2003). The model therefore treats E2 not as a weak accidental binder, but as a ligand capable of engaging AR and biasing it toward reduced stability and output under normal regulatory constraints, thereby providing a buffering mechanism that limits excessive androgen signaling during periods of androgen overdrive. In healthy adult males, circulating testosterone concentrations exceed estradiol by approximately 100-500 fold, and even in females androgen levels often surpass estradiol outside the peri-ovulatory window (Sipe and Van Voorhis 2007; Travison *et al*. 2014). Under these conditions, reciprocal androgen binding to ERs would be expected to overwhelm estrogen signaling fidelity.

**Figure 5.**
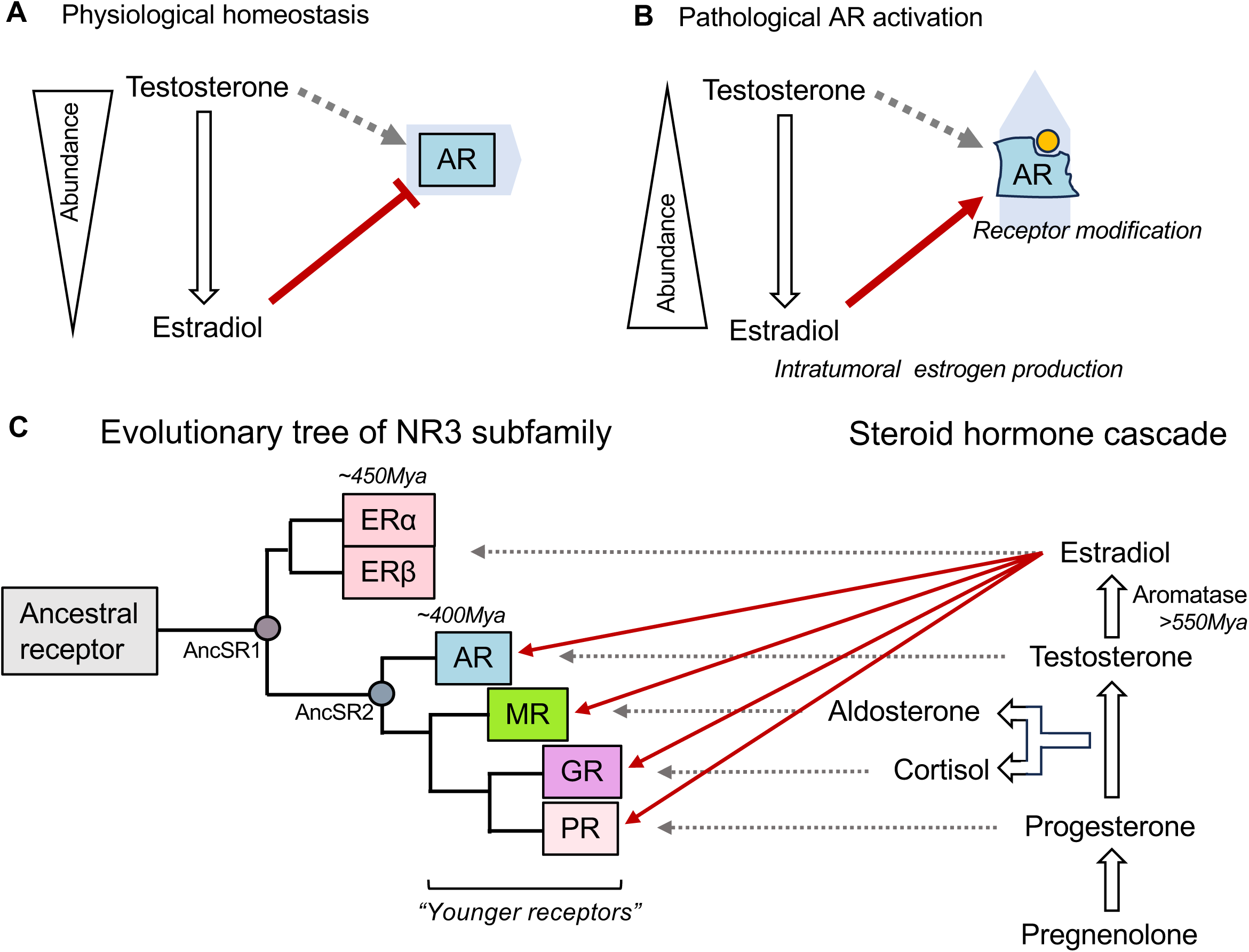
Evolutionary inheritance and pathological repurposing of estradiol-NR3 interactions. (A) Under normal endocrine conditions, circulating testosterone is present at substantially higher concentrations than estradiol, and AR preferentially responds to androgens. Estradiol nonetheless retains the ability to engage AR via an ancestral-compatible binding pocket, but this interaction biases AR toward reduced stability and output (red inhibitory bar), functioning as a buffering brake that limits excessive androgen signaling. (B) In pathological contexts such as castration-resistant prostate cancer, receptor modification (e.g. AR ligand-binding domain mutations or altered co-regulator environments) repurposes this ancestral estradiol compatibility. Estradiol binding is converted from a suppressive interaction into a stabilizing, transcriptionally productive signal (red arrow), enabling estradiol-driven AR activation despite low androgen availability. (C) Phylogenetic reconstruction of the NR3 subfamily illustrates that the earliest steroid receptor (AncSR1) was estrogen-responsive, reflecting estradiol as an ancient ligand predating diversification of steroid receptors. Subsequent duplication events produced ERα/β (∼450 Mya) and later AncSR2 (∼400 Mya), the common ancestor of AR, PR, GR, and MR. While ERs retained high specificity for aromatized estrogens, younger receptors progressively acquired responsiveness to non-aromatized steroids while retaining ancestral estradiol compatibility. The steroidogenic pathway proceeds from pregnenolone through progesterone and androgens to estradiol via aromatization (CYP19A1), which arose prior to modern NR3 diversification (>550 Mya). The persistence of estradiol affinity across younger NR3 receptors reflects inheritance of an ancestral binding architecture rather than de novo promiscuity. This evolutionary ordering explains the observed asymmetry: estradiol can engage AR, PR, GR, and MR, whereas upstream steroids do not reciprocally activate estrogen receptors. Together, this figure integrates physiological regulation, pathological rewiring, and evolutionary history to illustrate how estradiol, through a terminal biosynthetic product, acts as an ancestral ligand whose retained compatibility underlies asymmetric cross-reactivity and context-dependent modulation of NR3 receptor signaling.

In contrast, a pathological “maladaptive activation” state (Figure 5B) emerges when these constraints are relaxed or rewired. Prior work showed that the AR coactivator ARA70 can enable estradiol-driven AR transcriptional activation (Yeh *et al*. 1998), and that altered coactivator landscapes in advanced disease can lower the barrier for noncanonical ligand activation (Talukdar and Chatterji 2023). Clinically, ligand-binding domain mutations further expand AR permissiveness in a subset of castration-resistant prostate cancers (Brooke *et al*. 2008; Chung and Abboud 2022), including variants in which estradiol can stimulate AR target gene expression (Nakhla *et al*. 1997). This suggests that mutation tunes an inherited ligand compatibility that is already present, aligning with a broader principle in which disease exploits latent receptor properties embedded by evolutionary history rather than inventing new biochemical functions.

Together, these factors provide a mechanistic route by which estradiol’s basal compatibility with AR, which is normally used for buffering, can be co-opted into a growth-promoting activation mode during androgen deprivation.

### Estradiol engages a conserved AR pocket and mutations amplify, rather than invent, compatibility

A key mechanistic implication is that estradiol does not require a specialized or mutation-exclusive binding mode to engage AR. Instead, estradiol and testosterone occupy a shared canonical binding pocket in AR (Figure 2A), consistent with a conserved steroid-recognition architecture. This structural compatibility provides an explanation for why estradiol can bind AR with high predicted affinity without requiring extensive remodeling of the receptor. Importantly, modeling of the clinically relevant AR T877A mutation indicates only modest potentiation of the pre-existing affinity landscape across ligands, with estradiol remaining among the strongest predicted ligands (Figure 3A-C). This pattern is most consistent with “repurposing” of an ancestral estradiol-compatible interaction to stabilize and activate AR in permissive contexts, rather than creation of a novel binding mode. In other words, mutation tunes an inherited compatibility that is already present, aligning with the broader conclusion that disease exploits latent features embedded by evolutionary history. Such maladaptive receptor modifications may preferentially stabilize the active AR conformation and enhance transcriptional competence (Gim *et al*. 2021) rather than substantially increasing ligand-binding affinity itself.

### Evolutionary origin: estradiol as an ancient ligand retained across NR3 diversification

The directionality of cross-reactivity is naturally explained by the evolutionary history of NR3 receptors. Phylogenetic and functional reconstructions support that the earliest steroid receptor was estrogen-sensitive (Thornton 2001; Thornton *et al*. 2003), and that AR, PR, GR, and MR emerged later through duplication and divergence events in early vertebrates (Dehal and Boore 2005; Jaillon *et al*. 2004; Thornton 2001). Because these younger receptors inherited their ligand-binding scaffold from an estrogen-sensitive ancestor (Thornton 2001), estradiol compatibility is expected to persist even as specificity for non-aromatized ligands evolves.

Our ancestral receptor analysis (Figure 4) provides direct support for this evolutionary logic. AncSR1 is estrogen-responsive in predicted affinity, consistent with an ancestral estrogen-sensitive state, while AncSR2 shows a broadened affinity landscape including multiple non-aromatized steroids, consistent with an intermediate stage preceding full receptor specialization. Across the affinity heatmap (Figure 4D), estradiol maintains consistently high predicted affinity across ancestral and modern receptors, whereas other steroids show progressively diversified receptor preferences following NR3 lineage expansion. This pattern supports a “ligand exploitation” model. Estradiol, though biosynthetically downstream, acts as the evolutionarily older ligand, and younger receptors diversified to recognize upstream steroids while retaining ancestral sensitivity to estradiol (Thornton 2001).

Figure 5C summarizes the conceptual inversion revealed by this analysis. Although estradiol is a terminal product of the steroidogenic cascade, it predates strict specialization of modern NR3 receptors, and its structural features remain compatible with the conserved NR3 pocket. By contrast, ERα/β, which was already optimized for aromatized estrogen recognition, never evolved responsiveness to non-aromatized steroids, producing the asymmetric pattern observed across extant receptors.

### Estradiol as an end-product buffer in steroid physiology, and cancer as a trade-off exploit

The persistence of estradiol compatibility across NR3 receptors suggests physiological value. Steroid hormones share a common origin from cholesterol and form an interconvertible metabolic cascade. Under these conditions, end-products can serve as stabilizing feedback signals. We propose that estradiol functions as a substrate-coupled buffer that restrains activity of receptors sensing upstream intermediates (Figure 5A), providing a unidirectional control mechanism that limits runaway activation in androgenic, progestational, or corticoid axes. Conversely, reciprocal binding of abundant upstream steroids to ERs would risk signal corruption due to their higher circulating concentrations(Travison *et al*. 2014), making selection for ER insulation plausible.

Malignancies can exploit this architecture (Figure 5B). During androgen deprivation, tumors gain selective advantage by enabling estradiol to serve as a functional AR agonist through coactivator enrichment(Yeh *et al*. 1998), AR ligand-binding domain mutation(Brooke *et al*. 2008; De Bosscher *et al*. 2020; Nakhla *et al*. 1997; Talukdar and Chatterji 2023), and increased local estrogen production. Reports of aromatase expression in prostate cancer support the plausibility of intratumoral estrogen availability(Ellem *et al*. 2004), and this context provides a mechanism by which a physiological buffering ligand becomes a pathological driver in castration-resistant disease. This framework also rationalizes why aromatase inhibition may produce heterogeneous outcomes depending on whether estradiol is acting primarily as a brake (Figure 5A) or as a fuel source (Figure 5B) (Hoang *et al*. 2017; Wadosky and Koochekpour 2016).

### Extending the buffering concept across NR3: PR, GR, and MR as parallel substrates for estradiol engagement

Although AR provides a clinically salient example, the same architectural constraints likely apply broadly across NR3 receptors. Our affinity screen predicts high estradiol engagement of PR, GR, and MR as well, (Figure 1G-I) consistent with shared descent from an estrogen-sensitive ancestor (Thornton 2001). This supports a unifying principle: estradiol, as an aromatized end-product, can act as a system-level regulator of upstream steroid axes via direct receptor engagement (Figure 5C).

In PR biology, estrogen-progesterone antagonism is classically interpreted through receptor crosstalk and transcriptional network effects (Mohammed *et al*. 2015; Singhal *et al*. 2016), but direct E2-PR binding predicted here offers an additional mechanistic layer that could contribute to context-dependent progesterone antagonism. In GR physiology, estradiol is known to modulate stress and immune responses (Kalaitzidis and Gilmore 2005; Straub 2007), and direct receptor engagement may help explain estradiol-associated glucocorticoid resistance in some settings (Krishnan *et al*. 2001). In MR biology, estradiol-dependent modulation of MR-driven programs has been described in cardiovascular and renal contexts (Bienvenu *et al*. 2022; Metcalfe and Meldrum 2006), and direct E2-MR engagement could contribute to these effects alongside canonical ER-mediated mechanisms. Across these receptors, the same two-state logic may apply: a physiological buffering mode versus a pathological or permissive activation mode driven by altered cofactors, receptor variants, or local steroid environment.

### Testable predictions, limitations, and outlook

This framework generates testable predictions with translational relevance. Tumors that activate AR through estradiol under permissive states, such as those harboring promiscuous AR mutations including T877A or H874Y (Brooke *et al*. 2008; Elo *et al*. 1995; Nakhla *et al*. 1997), should be selectively vulnerable to interventions that disrupt E2-AR binding while preserving androgen-directed pharmacology. Because the activation state requires a permissive cofactor environment, reducing ARA70/NCOA4 or related coactivators is predicted to convert estradiol from an agonist back to a destabilizing or antagonistic ligand (Woodham *et al*. 2003; Yeh *et al*. 1998). Finally, tumors with elevated local estrogen production are predicted to show diminished responses to AR-only strategies and may exhibit state-dependent sensitivity to aromatase inhibition (de Ronde and de Jong 2011), consistent with heterogeneous clinical outcomes.

Several limitations should be acknowledged. Boltz-2 predictions provide a scalable affinity landscape but do not replace biochemical binding measurements or functional assays. The model has particular limitations for distinguishing between binding modes that may differ in their functional outcomes. In addition, predicted affinity alone does not specify transcriptional outcome, emphasizing the importance of the two-state model and the need for direct experimental validation in defined cofactor and mutation contexts. The ancestral receptor reconstructions, while based on robust phylogenetic methods, carry inherent uncertainty about precise functional states of ancient proteins.

### Conclusion

We define an asymmetric cross-reactivity within NR3 steroid receptors in which estradiol retains strong compatibility with later evolved receptors while reciprocal binding of upstream steroids to ERs is excluded. By integrating comprehensive affinity prediction, conserved pocket occupancy, mutation effects, and ancestral reconstruction, we propose that estradiol’s broad NR3 engagement reflects an evolutionarily retained buffering architecture that can be repurposed in disease. This conceptual framework provides a unifying explanation for estradiol’s paradoxical effects on AR signaling and motivates renewed attention to the E2-AR interface as a context-dependent therapeutic vulnerability.

## Declaration of interest

The authors declare that there is no conflict of interest that could be perceived as prejudicing the impartiality of the research reported.

## Acknowledgement

A.B.R. acknowledges funding from the Perelman School of Medicine, University of Pennsylvania, the Institute for Translational Medicine and Therapeutics (ITMAT) at the University of Pennsylvania. This work was supported also by NIH DP1DK126167 and R01GM139211.

## Author contributions

S.Y. performed the data analysis and wrote the manuscript.

A.B.R. designed the project and analyzed results, secured funding, and wrote the paper with contributions from the other authors. All authors have reviewed the final manuscript and agree on its interpretation.

## Supplementary files

**Supplementary File 1**

Predicted ligand-receptor affinity matrix for all human nuclear receptors and ligands analyzed, including IC₅₀ estimates and confidence scores.

## Notes

### Competing Interest Statement

The authors have declared no competing interest.

